# Reliance on Episodic vs. Procedural Systems in Decision-Making Depends on Individual Differences in Their Relative Neural Efficiency

**DOI:** 10.1101/2023.01.10.523458

**Authors:** Yuxue Yang, Catherine L. Sibert, Andrea Stocco

**Author notes:** Corresponding author: Andrea Stocco, Department of Psychology, Campus Box 351525, University of Washington, Seattle, WA 98195.

## Abstract

Experiential decision-making can be explained as a result of either memory-based or reinforcement-based processes. Here, for the first time, we show that individual preferences between a memory-based and a reinforcement-based strategy, even when the two are functionally equivalent in terms of expected payoff, are adaptively shaped by individual differences in resting-state brain connectivity between the corresponding brain regions. Using computational cognitive models to identify which mechanism was most likely used by each participant, we found that individuals with comparatively stronger connectivity between memory regions prefer a memory-based strategy, while individuals with comparatively stronger connectivity between sensorimotor and habit-formation regions preferentially rely on a reinforcement-based strategy. These results suggest that human decision-making is adaptive and sensitive to the neural costs associated with different strategies.

## Introduction

Two distinct frameworks have been proposed to explain how experience influences human behavior. According to one class of theories, previous decisions and their outcomes are stored as traces in episodic long-term memory, and decision-making is controlled by which memory is easier to retrieve (Gonzalez & Dutt, 2011; Stewart et al., 2006): we will refer to this as the declarative framework. The other class of theories suggests that decision-making based on experience is governed by the basic principles of reinforcement learning, in which the value of the outcome of each previous decision incrementally modifies an internal cached value associated with each option (Lee et al., 2012; Niv, 2009); when making a decision, the option with the highest expected value is chosen. We will refer to this as the procedural framework.

Despite making similar predictions about the outcome of decision-making processes (Chelian et al., 2015), these two mechanisms depend on different neural resources. In the *declarative* framework, decisions from experience depend on the brain circuits involved in the storage and retrieval of episodes in long-term memory, such as the medial temporal lobe (Knowlton et al., 1994), the lateral prefrontal cortex (Badre & Wagner, 2007), and the medial frontal and parietal regions associated with the default mode network (Raichle & Snyder, 2007; Sestieri et al., 2011). In the *procedural* framework, decisions from experience rely on brain circuits for implicit reward learning and habit formation, such as the basal ganglia (Foerde et al., 2006), the supplementary motor area (Nieuwenhuis et al., 2004), the cerebellum (Taylor & Ivry, 2014), and the specific sensorimotor regions involved in the task itself (Hill & Schneider, 2006).

Both systems are concurrently active at any given time (Foerde et al., 2006; Poldrack et al., 2001), so why should individuals choose to rely on one system over the other? One likely explanation is that optimal behavior in humans is inherently limited by processing capacity (Simon, 1957) and individuals must therefore adopt different procedures to maximize the outcome of a decision while consuming the minimum amount of cognitive resources (Gigerenzer, 2008; Payne et al., 1988, 1993). Because the cost of a decision also depends on the neural resources it consumes, individuals with different neural characteristics would make decisions in different ways. This paper puts forward the hypothesis that individuals rely on *declarative* or *procedural* processes based on the relative neural efficiency of their corresponding brain circuits.

A common indicator of neural efficiency within a circuit is their functional resting-state connectivity; that is, the degree of correlation between the spontaneous neural activities of different regions in that circuit (Shen et al., 2017). Higher correlations at rest reflect tighter coupling of neural dynamics and greater exchange and integration of communication between regions (van den Heuvel & Hulshoff Pol, 2010). Conversely, lower correlations could be interpreted as reduced information exchange or noisier communication channels between regions. All things being equal, an adaptive decision maker (Payne, Bettman, and Johnson, 1993) should preferentially rely on the most precise and efficient system for decision making. Thus, relatively higher functional connectivity within the *declarative* or *procedural* circuit should predict greater reliance on the corresponding system when making decisions.

An important obstacle in testing our hypothesis is that the choice between the two systems does not depend solely on their mental and neural costs but also on their relative effectiveness in a given task. For example, it has been argued that the *procedural* system would be preferred when decisions are probabilistic and the stimuli are difficult to verbalize (Frank et al., 2004; Knowlton et al., 1994; Poldrack et al., 2001). If, for some reason, one system is better suited for a given task than the other, an individual preference based on neural efficiency would be overridden by the effectiveness of the alternate system. Investigating the relationship between brain connectivity and preferred decision-making processes requires a task where the *procedural* system is equally as effective as the *declarative* system strategy. One such task is the Incentive Processing Task (IPT: Delgado et al., 2000), which guarantees that either system yields the same expected payoff. In this task, participants repeatedly guess whether a hidden number is greater or smaller than five by pressing one of two buttons, and receive monetary feedback for correct guesses. Once the choice is made, the number is revealed, and feedback is provided (Figure 1A). Unbeknownst to participants, the number is chosen after their decision is made and follows a predefined feedback schedule. Under these conditions, preferences for either the *declarative* or *procedural* decision-making systems should depend only on the neural costs for each individual participant. Because the number of wins and losses is fixed and predefined, behavioral data from the IPT cannot be analyzed using accuracy measures. Instead, individual behavioral differences can be measured by computing the probability of choosing a different option (i.e., from “smaller” to “greater” or vice versa) after receiving feedback from the previous trial, referred to as the shift probability. Shift probabilities were computed for the different types of feedback (“win,” “lose” or “neutral” when the number is exactly five) and the two different types of experimental blocks that were used in the task (“Mostly Lose,” in which 6 loss trials were pseudo-randomly interleaved with either 1 neutral and 1 reward trial, 2 neutral trials, or 2 reward trials, and “Mostly Win”, in which 6 reward trials were pseudo-randomly interleaved with either 1 neutral and 1 loss trial, 2 neutral trials, or 2 loss trials). Under these conditions, individual preferences for *declarative* vs. *procedural*-based decision-making should maximally reflect the underlying efficiency of the corresponding circuit.

**Figure 1.**
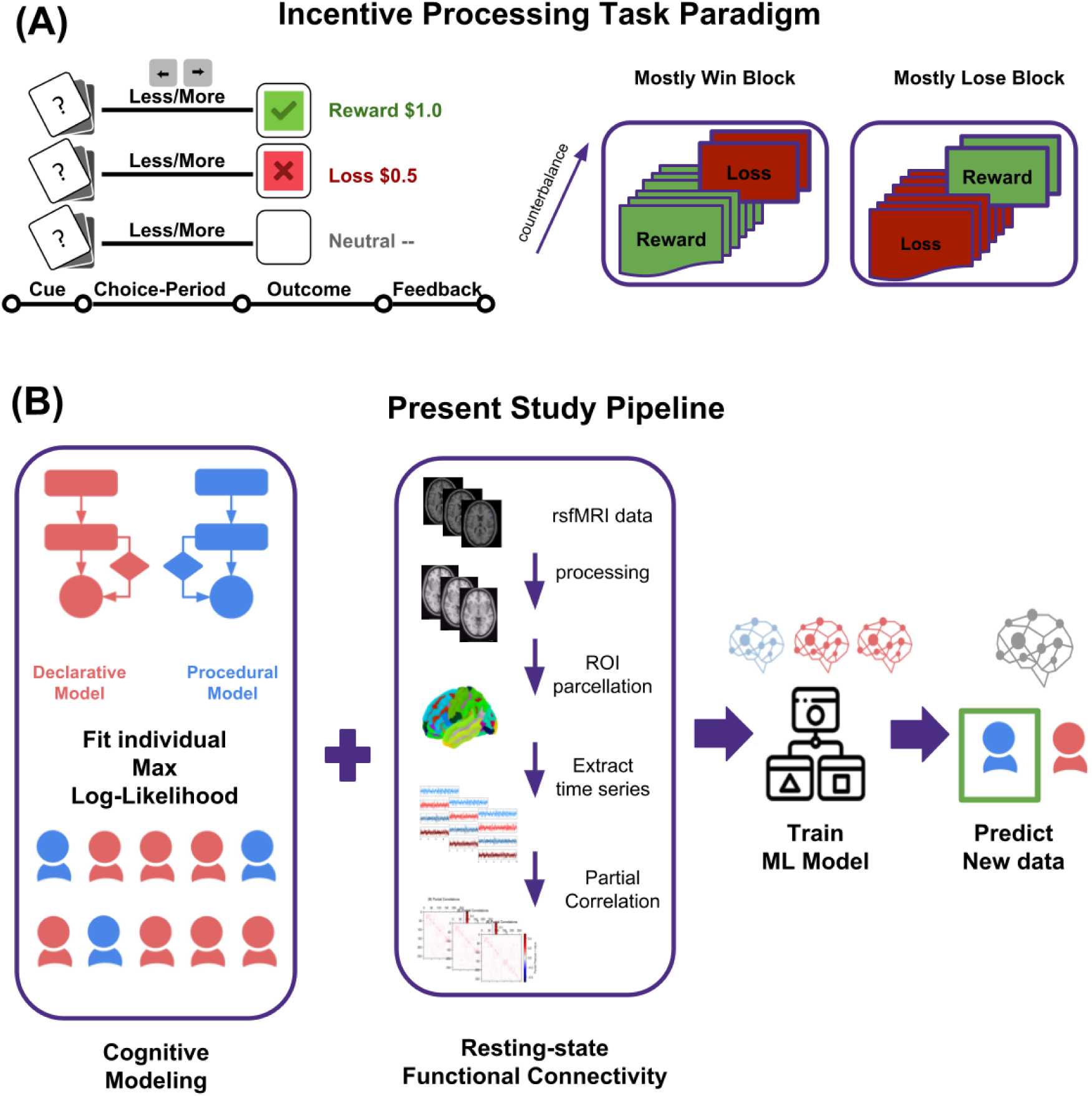
(A) Incentive processing task paradigm. (B) The pipeline of the present study. First, each subject’s behavioral data are fit by two distinct cognitive models, *declarative* vs. *procedural* model through maximum log-likelihood. Second, each subject’s neuroimaging data are processed to construct resting-state functional connectivity matrices. Lastly, machine learning models are trained to predict an unseen subject’s decision-making process from the neuroimaging data.

To test this hypothesis, we analyzed a subset of 200 participants from the Human Connectome Project (Van Essen et al., 2013) for whom neural and behavioral measures from the IPT as well as resting-state fMRI data were available. Figure 1B illustrates the methodology of this study. To determine whether a participant relied on *procedural* or *declarative* mechanisms, each participant’s behavior was fitted to two parametrized computational models, one implementing *declarative* decision-making and one using the *procedural* system.

The episodic control model is based on the memory model of Anderson and Schooler (Anderson & Schooler, 1991), which follows the multiple trace theory (Nadel et al., 2000) and has an established interpretation in terms of brain circuitry and anatomy (Stocco et al., 2023). In this model, the outcome of every decision is stored as an episodic memory. After encoding a memory, without any intervention, that memory starts to be forgotten, and the availability of each memory trace decays according to a power law (Newell & Rosenbloom, 1981). Multiple experiences of the same decision-outcome pair, however, result in the accumulation of traces of the same memory, increasing the *activation* of that memory. Formally, the activation of a memory *m* at time *t* reflects the balance between these two effects, and is given by the equation:

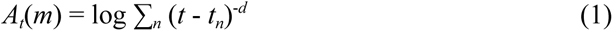

In this equation, *t_n_* is the time at which the same decision/outcome pair has been experienced for the *n*-th time and *d* is an individual-specific forgetting rate (Sense et al., 2016; Zhou et al., 2021). A decision is guided by the most active memory of past interactions. If the retrieved memory is of a “win” outcome, the model repeats its associated decision, (e.g., choosing “more” if the “win” was associated with a “more” choice). If the memory is of an associated loss, the model executes the *opposite* action (e.g., choosing “less” if the loss was associated with a “more” choice). The probability that a given memory *m* will be retrieved is controlled by a Boltzmann function with noise temperature *ν*:

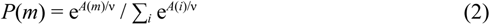

The *procedural* model is based on the reinforcement learning framework, which has a similarly established interpretation in terms of neuroanatomy (Niv, 2009). In this model, the two possible decisions (“more” or “less”) are represented as two actions associated with the cue (in this paradigm, this is the presentation of the two cards). Each action *a* has an associated value *V*(*a*) that is updated over time based on the difference between the expected and the actual outcome (Rescorla et al., 1972). For example, receiving a “win” feedback after making a “more” choice will increase the value of the “more” action, *V*(*more*), making it more likely to be selected in future choices, while receiving “loss” feedback will decrease the value of the “more” action, making it less likely to be selected in future choices. Formally, after the *n*-th decision, the value *V_n_*(*a*) of an action is updated using the temporal difference method (Sutton, 1988):

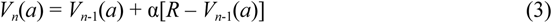

In Equation 3, *R* is the reward associated with an outcome, (1 for “win” and –0.5 for “lose”, corresponding to the dollar amount won by participants), and α is the learning rate. The probability that an action *a* will be selected is controlled by a Boltzmann function with noise temperature τ:

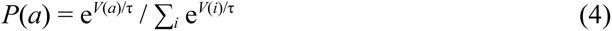

Thus, each model depends on one learning parameter (*d* or α) and one noise parameter (ν or τ).

Both models were fit to each participant using maximum likelihood estimation (Myung, 2003), i.e., by fitting their parameters to maximize the probability of generating the specific series of decisions made by that participant (see Methods). Each participant was then assigned to either the *declarative* or *procedural* group based on which version of the model had the greatest likelihood of producing their observed behavioral data.

Although indistinguishable in most cases, the two models behave differently in the IPT. Because both models adaptively adjust their behavior after a new feedback is received, both models predict that the probability of switching response would be greater after a loss than after a win. The declarative model alone, however, also exhibits the effect of *recency*, that is, the fact that the more recent outcomes have greater impact while older outcomes fade away. This implies that the effect of trial feedback on the declarative model should be magnified, with a recent loss resulting in a greater probability of switching and a recent win resulting in a smaller probability of switching between responses. To test this intuition, we analyzed data from 1,000 simulated runs of each model, drawing different values for the model parameters from uniform distributions over representative ranges of their values. To choose these representative ranges, we considered that recommended values of α are typically around 0.1 and recommended values of *d* are typically around 0.5. Thus, we explored values of 0.0 < α < 0.5; 0.25 < *d* < 0.75, and values of temperature parameters *ν* and *τ* between 0 and 1. The results for these forward simulations are shown in Figure 2:

**Figure. 2:**
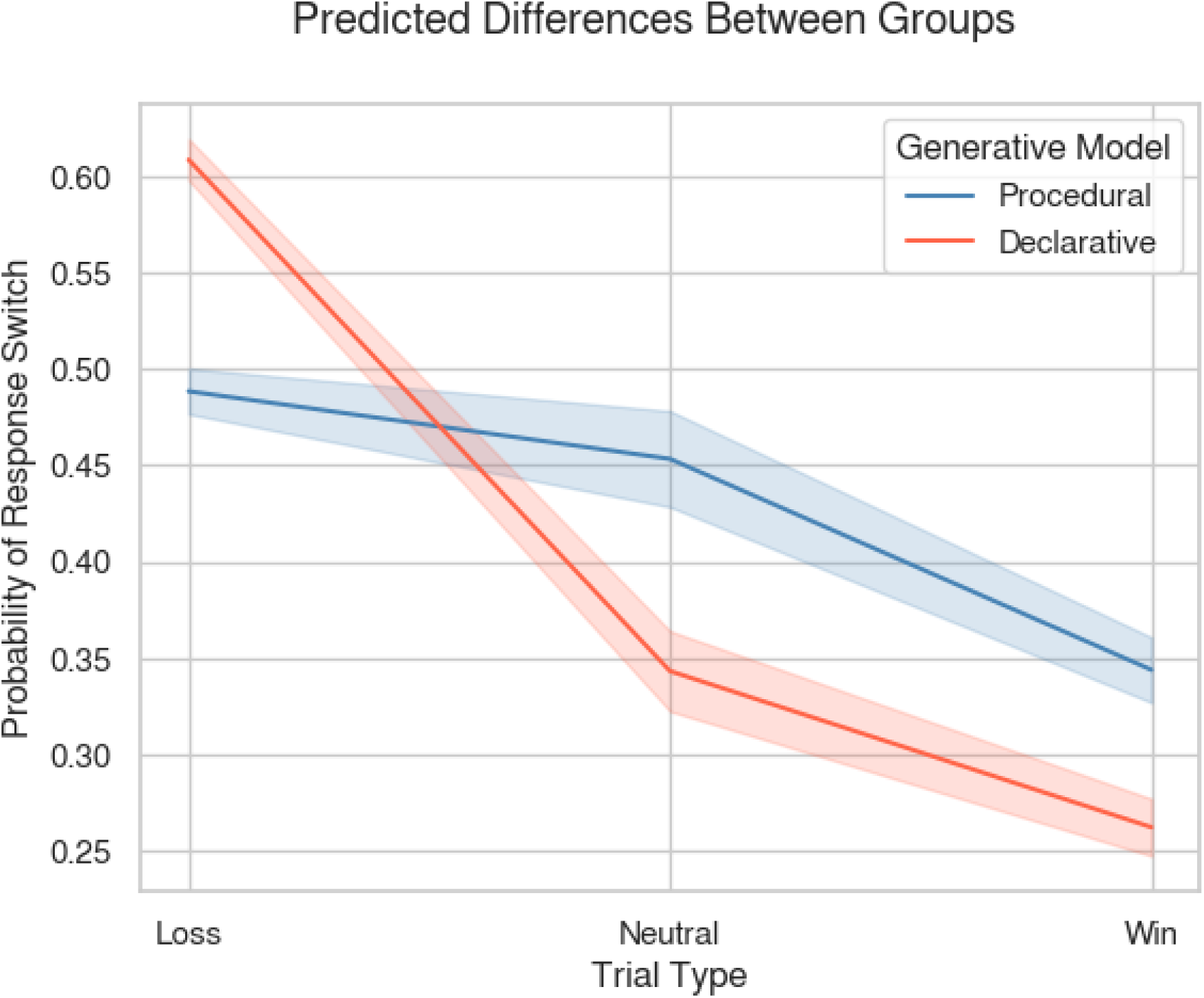
Predictions from simulations of the two models. Each model was run 1,000 times with randomly initialized parameters drawn from a uniform distribution over representative parameter ranges (0.0 < α < 0.5; 0.25 < *d* < 0.75, 0 < *ν* < 1, and 0 < *τ* < 1; See main text).

As the figure shows, the declarative model shows a greater sensitivity to the trial type (e.g., greater changes in switch probabilities from losses to neutral to win trials) than the procedural model, resulting in a model by trial type interaction.

Thus, at the behavioral level, we expect individuals best fit by the declarative model to show a corresponding greater effect of trial type than those best fit by the procedural model, resulting in a similar model by trial type interaction.

And because the two mechanisms rely on different neural substrates, we expect that regional differences in BOLD activation would reflect the neural cost associated with each strategy. Therefore, we expect individuals best fitted by *declarative* models to show greater activity in regions associated with episodic encoding, such as the hippocampus, and retrieval, such as the ventral prefrontal cortex and default mode network (Stocco et al., 2023). Correspondingly, we expect individuals best fit by the *procedural* model to display greater activity in the circuits associated with habit formation, such as the basal ganglia, the cerebellum, and the visuo-motor regions involved in interacting with the stimuli (Ashby, Turner, & Hovitz, 2010). Finally, we expect that the type of decision-making processes used will depend on the patterns of functional connectivity at rest. To test this latter prediction, a classifier was trained to predict whether a participant would be best fit by a *declarative* or *procedural* model from the same individual’s functional connectivity data. We expect that functional connections that successfully predict the reliance on the *declarative* mechanism for decision-making would be found between memory encoding and retrieval regions. Conversely, we expected that functional connections predictive of reliance on the *procedural* mechanism reward-based learning would be found in sensorimotor cortices and the basal ganglia circuit.

## Methods

### Materials

This study analyzed both behavioral and neuroimaging data obtained from a subset of the Human Connectome Project (HCP) dataset (Van Essen et al., 2013). Data were provided by the Human Connectome Project, WU-Minn Consortium (Principal Investigators: David Van Essen and Kamil Ugurbil; 1U54MH091657) funded by the 16 NIH Institutes and Centers that support the NIH Blueprint for Neuroscience Research and by the McDonnell Center for Systems Neuroscience at Washington University. A total of 199 participants (111 females, 85 males, and 3 did not disclose) who completed both sessions of the task-based fMRI gambling game were included in this study and fit by the two models. By excluding 23 participants who missed resting-state fMRI scanning, a total of 176 participants were fit by a predictive LASSO model. All participants were healthy adults with no neurodevelopmental or neuropsychiatric disorders. The experimental protocol, subject recruitment procedures, and consent to share de-identified information were approved by the Institutional Review Board at Washington University.

### The Incentive Processing Task in the HCP

This incentive decision-making task was adapted from the gambling paradigm developed by Delgado and colleagues (Delgado et al., 2000). Participants were asked to guess if the number on a mystery card (represented by a “?”, and ranging from 1 to 9) was more or less than 5. After making a guess, participants were given feedback, which could take one of three forms, “Win” (a green up arrow and $1), “Loss” (a red down arrow and −$0.50), or “Neutral” (a gray double-headed arrow and the number 5). The feedback did not depend on the subject’s response, but was determined in advance; the sequence of pre-defined feedback was identical for all participants. The task was presented in two runs, each of which contains 32 trials divided into four blocks. Blocks could be “Mostly Lose” (6 loss trials pseudo-randomly interleaved with either 1 neutral and 1 reward trial, 2 neutral trials, or 2 reward trials) or “Mostly Win” (6 win trials pseudo-randomly interleaved with either 1 neutral and 1 loss trial, 2 neutral trials, or 2 loss trials). In each of the two runs, there were two “Mostly Win” and two “Mostly Lose” blocks, interleaved with 4 fixation blocks (15 seconds each). All participants received money as compensation for completing the task, and the amount of reward is standard across subjects.

### fMRI Data Processing and Analysis

This study employed the “minimally preprocessed” version of resting-state fMRI data and incentive processing task fMRI data, which has already undergone a minimal number of standard preprocessing steps including artifact removal, motion correction, normalization, and registration to the standard MNI ICBM152 template. Additional preprocessing steps were performed using the AFNI software (Cox, 1996, 2012), including despiking, spatial smoothing with an isotropic Gaussian 3D filter FWHM of 8 mm, and removal of linear components related to the six motion parameters and their first-order derivatives.

Functional connectivity measures were constructed from the HCP resting-state data using Power et al.’s whole-brain parcellation (Power et al., 2011). This parcellation was used to construct a 264 Region of Interest (ROI) functional atlas, with each ROI containing 81 voxels. This parcellation atlas is defined in the MNI space and was applied to all participants in the HCP dataset. The extraction of the time series and calculation of the connectivity matrices was performed using R (R Core Team, 2020) and Python. Pearson correlation coefficients and partial correlation coefficients between the time series of each brain region were calculated for each participant, resulting in a 264 × 264 symmetric connectivity matrix for each session for each subject. The average correlation coefficients across subjects were calculated by first transforming each r value into a Z-value, and then retransforming the average Z value back into an equivalent *r* value using the hyperbolic tangent transformation (Silver & Dunlap, 1987).

For task-based fMRI data analysis, we specified the first-level analysis model and estimated the parameters corresponding to the difference between “Mostly Win” and “Mostly Lose” blocks, as in (Barch et al., 2013). The resulting contrast maps for each subject were then used in a second-level weighted t-test between-declarative and procedural groups. The test was implemented using AFNI’s 3dttest++ software, and its weights corresponded to the absolute difference in log-likelihood between the best-fitting declarative and best-fitting procedural model. This way, the contribution of each observation was proportionally scaled to the evidence favoring each participant’s assignment to their groups. The statistical significance level was set at a significance level of *q* < 0.01 corrected for multiple comparisons using a False Discovery Rate (FDR (Benjamini & Hochberg, 1995)) procedure.

### Computational Models

#### Episodic Memory Model

The episodic memory model relies on the declarative model of Anderson and Schooler (Anderson & Schooler, 1991), currently implemented as part of the influential ACT-R cognitive architecture (Anderson, 2009). When presented with a mystery card, the model retrieves a previous episodic memory in which the number was guessed successfully. After being presented with feedback, the model encodes a new memory associating the action with its outcome. Memories are retrieved based on their activation, a noisy quantity that depends on the frequency and recency with which the decision-outcome episodes have been experienced (Eq. 1).

#### Procedural Model

By contrast, the procedural model represents the possible actions of the decision-making processes as competing stimulus-response rules, and reinforcement learning is used to increase the use of the rule that leads to the best outcomes. Instead of encoding each trial as a memory of action and associated feedback, the model has two competing actions that implement the “More” and “Less” decisions. When presented with the mystery card, the model chooses one of the rules to execute based on its expected value. Initially, both rules have equal value, and one will be chosen at random. After each decision, the model is presented with a “Win”, “Lose”, or “Neutral” response, and this feedback is encoded as the reward term in the reinforcement learning equation (Eq. 2; +1 for a “Win” result, –0.5 for a “Lose” result, and 0 for a “Neutral” result). Positive rewards will encourage the model to repeat the associated action, while a sequence of losses will decrease the value of an action and encourage the selection of the alternate action.

### Model Fitting

Model parameters were fit to maximize their likelihoods given the data. The likelihood of a model *M* with parameters *θ* given a vector of data **x**, indicated as *L*(*M*, *θ* | **x**), is the probability of observing the data, given the model and its parameter, that is *L*(*M*, *θ* | **x**) = *P*(**x** | *M*, *θ*). The data **x** consists of a vector of decisions *x*_1_, *x*_2_, …, *x_N_*:

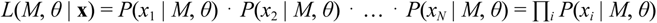

Since the product of conditional probabilities becomes vanishingly small, it is common to use *log*-likelihood:

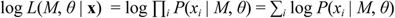

Note that, because both models are *memory* models, the probability of decision *x_i_* depends on the *history* of previous decisions *x_i_*_-1_, *x_i_*_-2_, …, *x*_1_ and their outcomes, i.e. *P*(*x_i_* | *M*, *θ*) = *P*(*x_i_*| *M*, *θ, x_i_*_-1_, *x_i_*_-2_, …, *x*_1_). In both models, the history of previous decisions is implicitly represented in their respective memory representations, that is, the activations of episodic memories and the values of actions. Thus, the probabilities of generating a response *x_i_* can be derived by Equations 2 and 4, provided that the history of previous decisions and outcomes is recorded in the corresponding activations and values.

To do so, each model is run forward, first calculating the probability of producing the participant’s response in the current trials; next updating the cumulative log-likelihood; and finally adjusting the activation of episodic traces and the value of the decision based on Equations 1 and 3 before moving to the next trial. For each participant, the model simulates the exact series of 64 trials as they experienced them, in the exact order and with the same inter-trial and inter-block intervals.

In order to evaluate the goodness-of-fit for individual fitting, we estimated maximum Log-Likelihood across the parameter space, i.e. we selected the value of parameters *θ*\* such that *θ*\* = argmax*_θ_ L*(*M*, *θ* | **x**). Because no closed-form solutions exist to derive the maximum likelihood of Equations 1 and 3, the best-fitting parameter values were identified using Powell’s optimization algorithm (Powell, 1964) as implemented in Python’s SciPy package (Virtanen et al., 2020). This method was chosen because it does not require explicit derivatives and allows us to specify meaningful bounds for parameter values. All parameters were constrained within the bounds [0, 2]. This choice was made to capture potentially unusual, but possible, decision-making behavior that lies beyond what is allowed by typical parameter ranges. In particular, given the frustrating nature of the task, we expected that some individuals would exhibit abnormally high sensitivities to the most recent feedback (captured by high values of *d* and α) or extensive exploration behavior (captured by high values of the noise parameters).

The starting parameter values for the algorithms were α = 0.1, τ = 0.1 for the procedural model, and *d* = 0.5, ν = 0.1 for the declarative model. As noted above, the starting values of α and *d* were chosen as the most common values found in their respective modeling frameworks; they are, for example, also the default values of the corresponding parameters in ACT-R.

### Supervised Classification Model

To explore if individuals’ behavioral differences between declarative and procedural strategies are indicated by an individual’s underlying brain structure, we trained three of the most commonly used supervised classification models (the Logistic regression model, the Decision Tree model, and the Random Forest model), using resting-state functional connectivity as the input variable, and predicted the probability of a participant being labeled as either preferring the declarative or procedural strategy. Considering the equally high accuracy (> 0.8) among these classification models, we are confident in saying that our Machine Learning models work very well in predicting the strategy selection from an individual’s resting state neuroimaging data.

In addition to the predictive power of machine learning models, another very important dimension we need to carefully consider in Machine Learning related research is the model’s interpretability. While some ML models are excellent at predicting outcome variables, as complexity grows exponentially (as in Deep neural networks), they become a black box that is fundamentally difficult to interpret. Therefore, choosing an appropriate ML model with reasonable predictability and interpretability is critical. We chose the logistic regression model because it is a simple and powerful binary classification model that has been widely used in many fields of research, and leads itself naturally to predicting the likelihood of a choice being made. Rather than fitting continuous data as a straight line, the logistic regression model uses the logistic function to transform numerical values into probability values between 0 and 1. Specifically, the output value *x* of a linear model is transformed into a corresponding probability *P*(*x*) by the function:

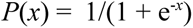

In order to handle an imbalanced dataset with unequal target labels, upsampling was applied by randomly resampling data from the minority class–in this case, the declarative group. We also applied individualized weights to each training sample, which is the absolute difference of maximum log-likelihood between two models. Specifically, for subjects who are better distinguished by either of the two models (declarative vs. procedural), we increased the weights of these data points in later ML model training, while for those who had a very close fit maximum log-likelihood between two models, the training procedure was less reliant on these samples. Having 69,696 (264 ROI × 264 ROI) connections, we want to select only the most important connections contributing to the prediction, therefore, LASSO regularization was applied to the Logistic Model. LASSO is a machine learning regression analysis technique that performs both variable selection and regularization in order to improve the prediction accuracy and interpretability of the computational model. It can reduce model complexity by penalizing large numbers of coefficients and also prevents overfitting which may result from a simple linear regression analysis. In LASSO, the best values associated with each regressor are chosen as to minimize a loss function that includes the ordinary squared errors as well as a penalty term for each regressor’s β weight:

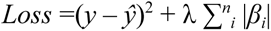

where the hyperparameter λ controls the degree of penalty: for greater values of λ, more coefficients are forced to become 0.

To account for the large disparity between the number of participants and the number of predictors, we performed a grid search cross-validation using the *glmnet* package in R (Friedman et al., 2010) to determine the best value for the fit hyper-parameter λ. To alleviate the potential problems of small sample size in neuroimaging studies, Vabalas and colleagues (Vabalas et al., 2019) used Nested CV approaches to produce robust and unbiased model performance regardless of sample size. In this approach, the validation follows two loops: an *outer* loop, in which the data is first split into training and testing set, and an *inner loop*, in which the training data is repeatedly split in sub-training and sub-testing data using cross-validation. Following their suggestions, we fit the model with *n*-iteration nested cross-validation (*n* = 200) to determine the optimal hyperparameter λ. For each outer loop iteration, the dataset was randomly split into training and testing. To enhance the model’s learning, we adopted 3:1 as training/testing split ratio. For the inner loop, we adopted the *k*-folds cross-validation (*k* = 20) method: the training dataset was randomly split into 20 folds and trained on 19 folds of samples. The prediction was made on the remaining one fold of samples to obtain the best hyperparameter value λ_k_, which gave the lowest classification error. With each best lambda λ_k_, the model was restrained and made predictions on testing datasets. Then the process was repeated *k* times. To guarantee maximum generalizability, we chose the median of the lambdas obtained in the outer loop (λ_nested_). This approach allowed us to obtain a reliable and unbiased assessment of the LASSO model’s performance while accounting for the variability in the data and the selection of hyperparameters.

The mean accuracy score, true positive rate (TPR), true negative rate (TNR), false positive rate (FPR), and false negative rate (FNR) were calculated across all folds to evaluate the overall performance of the model, taking sample weights into account. By definition, the receiver operating characteristic curve (ROC) demonstrates the performance of a classification model by plotting the relationship between TPR vs. FPR at different classification thresholds.As a measure of the model’s overall accuracy, we calculated the AUC (Area under the curve), which is one of the most important metrics for evaluating a classification model’s performance; as the AUC of a model approaches 1, the model approximates an ideal, perfect classifier.

## Results

### Reliability of Model Identification

Because our results depend on correctly identifying the model that best represents each participant, a model recovery procedure (Wilson & Collins, 2019) was run to ensure that our identification process was reliable. In this procedure, simulated participant data was generated by instantiating a model with random parameter values (uniformly drawn from within their boundaries; see Methods); the model-fitting procedure was then applied, treating the simulated data as participant data, and the accuracy to which the original model could be correctly identified was recorded (see Methods). Figure 3 provides the confusion matrix for the results of the procedure.

**Figure 3.**
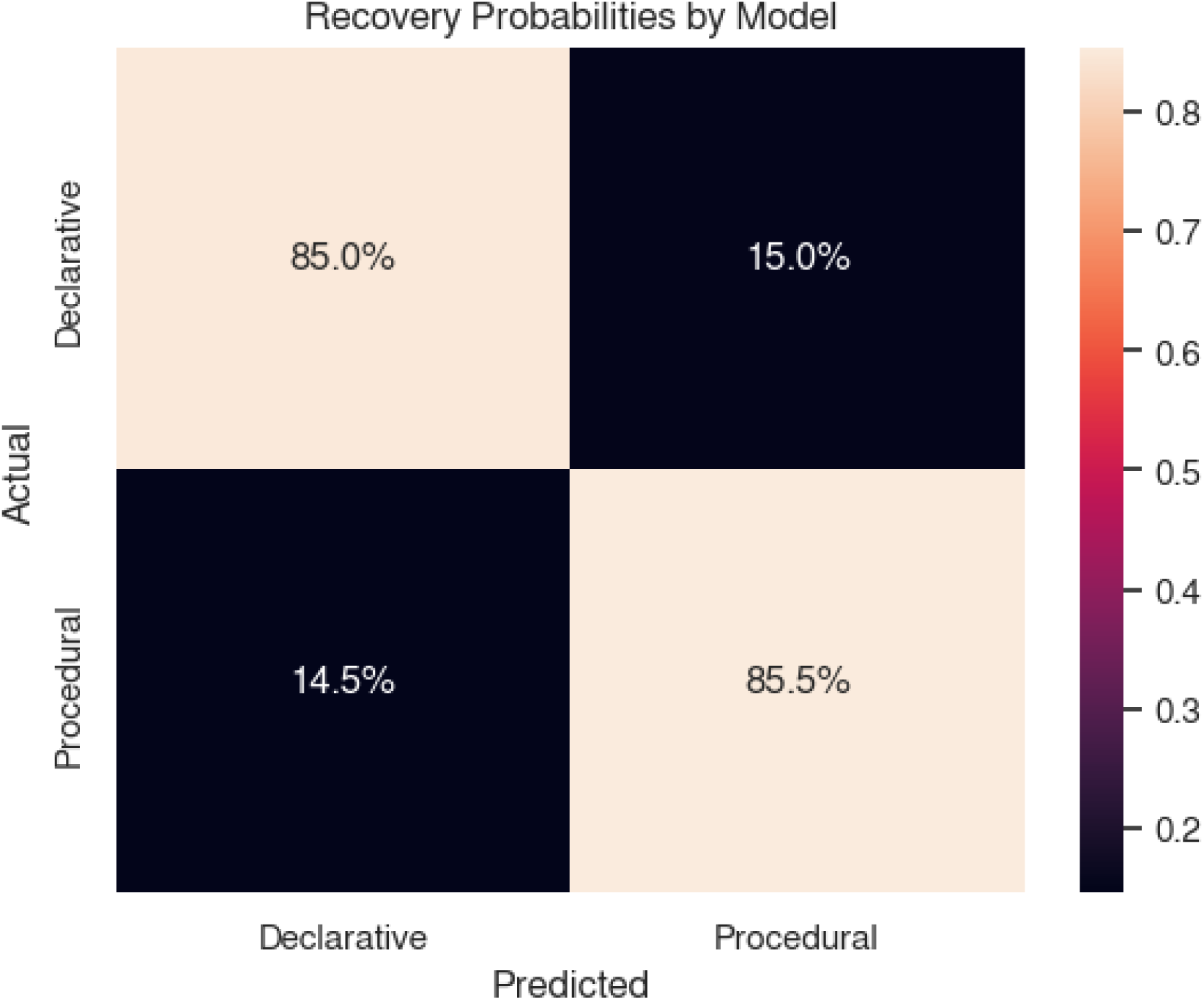
Confusion matrix showing the probability of correct model identification from 40,000 simulated runs. See Methods for details.

For both models, the recovery rate is approximately 85%. To account for errors in model assignment, in all of the following analyses, observations from a given participant are weighted by the difference in log-likelihood between the fit of the two models – that is, the stronger the evidence is in favor of one model, the greater the weight associated with the corresponding observation.

A second analysis was carried out to test whether each model’s *parameters* (i.e., *d* and *ν* for the declarative model and α and *τ* for the procedural one) could also be correctly identified. This was done to make sure that any inferences about individual differences in the relative efficiency of the two systems could be grounded on reliable differences in parameters. Figure 4 illustrates the results of these simulations. All four parameters could be correctly identified from the simulated data, with correlations ranging from *r* = 0.68 (procedural temperature τ, *p* < 0.001) to *r* = 0.88 (procedural learning rate α, *p* < 0.001).

**Figure 4.**
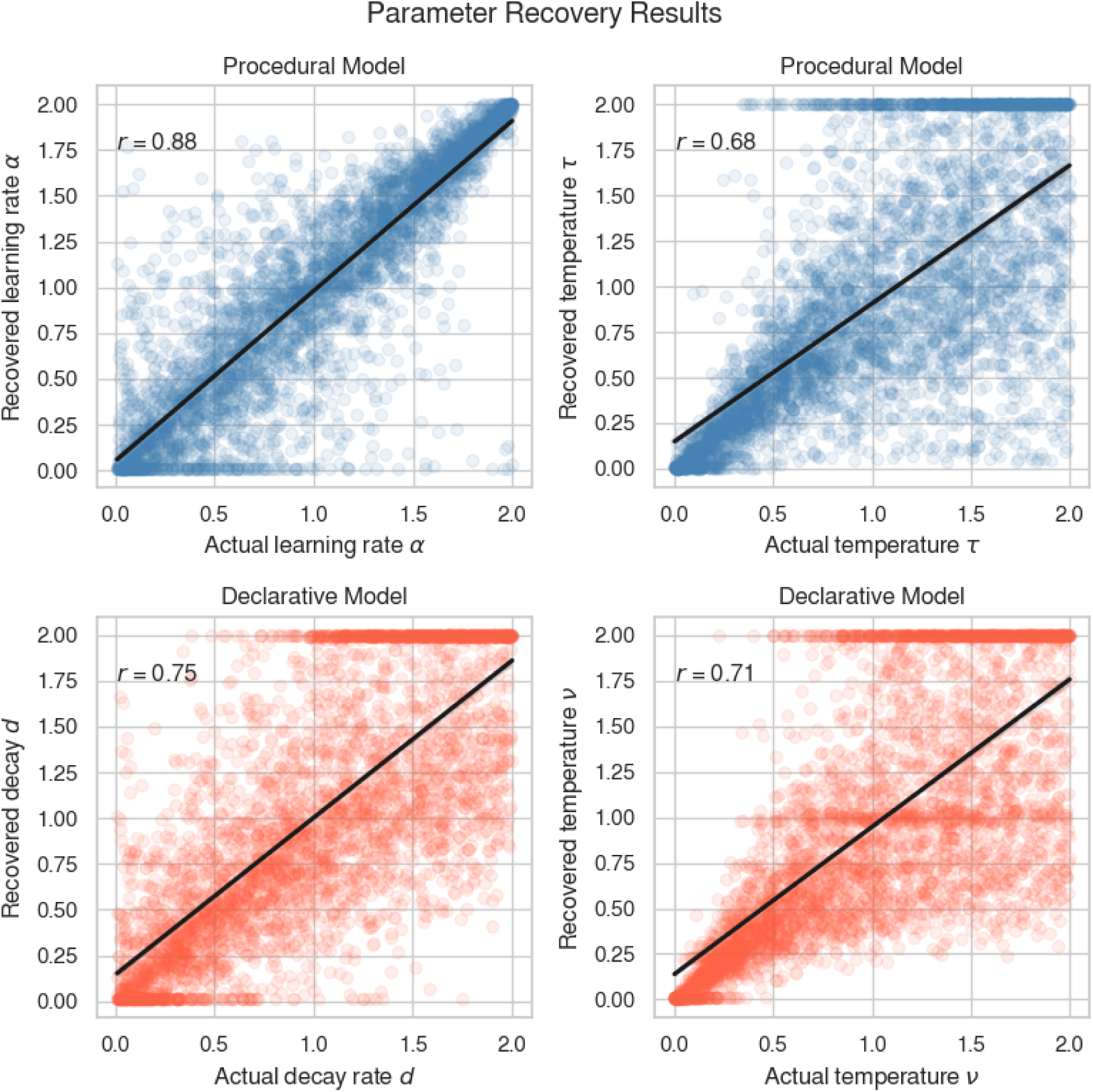
Results of the parameter recovery analysis for the individual parameters of the procedural (blue, top row) and the declarative (red, bottom row) models from 40,000 simulated runs. See Methods for details.

### Decision-Making Process Identification

Having examined the reliability of the models, we then proceeded with categorizing participants based on which model fit them best. A total of 199 participants were fit by the two models. Of these, 35 (17%) were best fit by the declarative model and thus were included in the declarative group.

The remaining 164 individuals were best fit by the procedural model and included in the procedural group. Figure 5 illustrates the distributions of the log-likelihood differences (Procedural - Declarative) across participants. Individuals for which the difference was < 0 were categorized as procedural, and those whose difference was > 0 were categorized as declarative.

**Figure 5.**
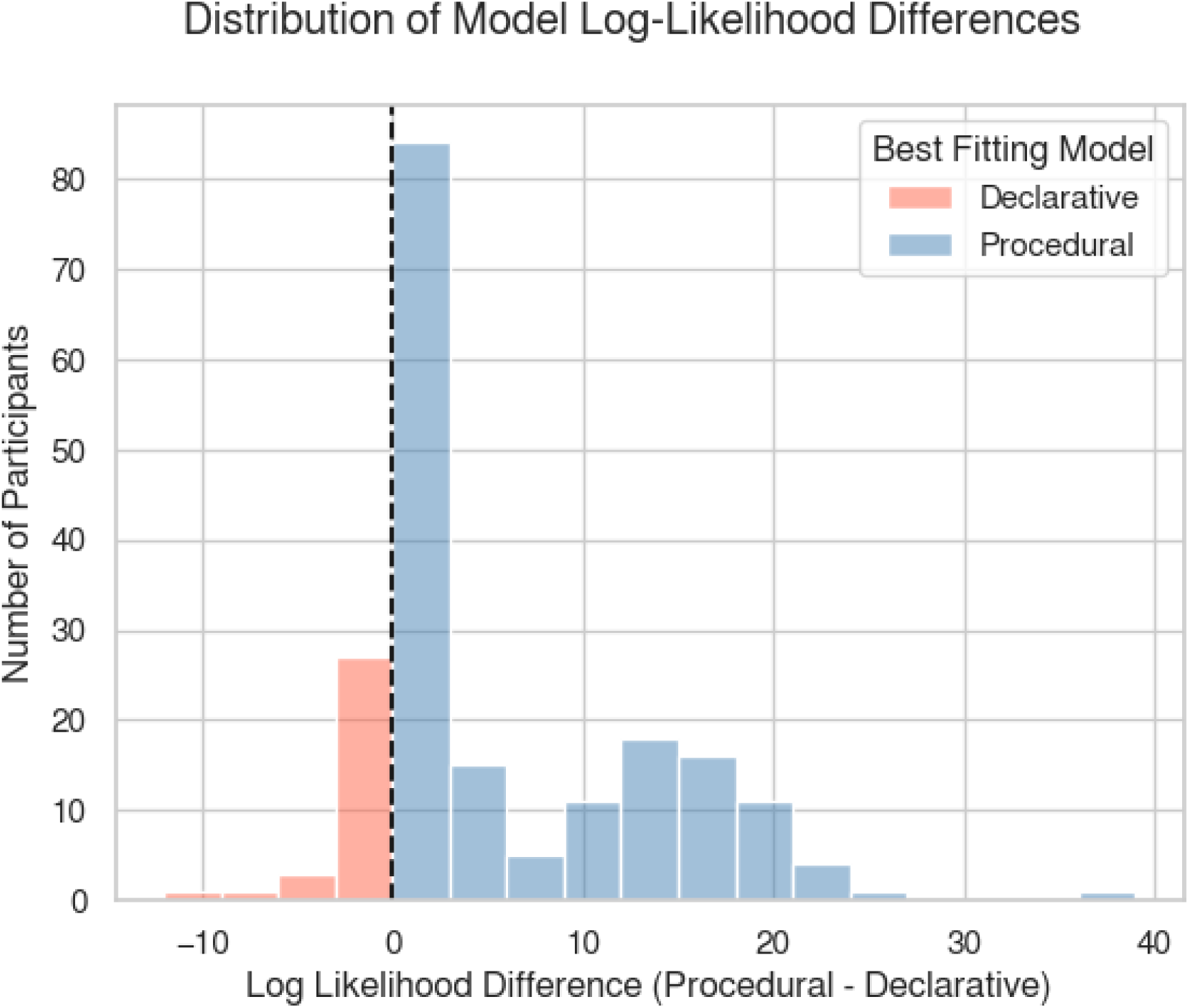
The distributions of the log-likelihood differences (Procedural - Declarative) across participants. As expected, the distribution is bimodal. Red bars represent the count of participants who are best fit by the procedural model, whereas blue bars represent the count of participants who are best fit by the declarative model.

In agreement with our hypothesis that individuals would preferentially perform the task using one of two mechanisms, the distribution of log-likelihoods is bimodal; that is, it has two peaks that likely correspond to the clusters of individuals best fit by the declarative and the procedural model. Note, however, that the leftmost peak is centered slightly to the right of the zero. This could be explained by the declarative model having inherently more difficulty fitting the data; this could happen, for example, if its equations do not capture the computations of human memory as well as the procedural model captures human reinforcement learning.

One possible solution to this problem would be to choose a decision criterion situated between the two peaks. In this study, we followed the alternative solution to keep the theoretically-motivated threshold point at zero, but to run all of the following analysis by weighting the contribution of each participant in proportion to the log-likelihood of their model identification. This second solution offers the advantage of maintaining a theoretically motivated criterion (that is, log *L* = 0) while also automatically accounting for the uncertainty in identifying certain participants (those whose log *L* values hover around zero).

We then visually inspected the distribution of parameter values of the best-fitting models for the two groups (Figure 6) to make sure that they conform to the values most typically used in the literature. The distributions conform to our expectations, with a subset of individuals exhibiting high decision noise and mean values of the decay and learning rates close to the established values for the corresponding models (mean α = 0.19 and a *d* = 0.64).

**Figure 6.**
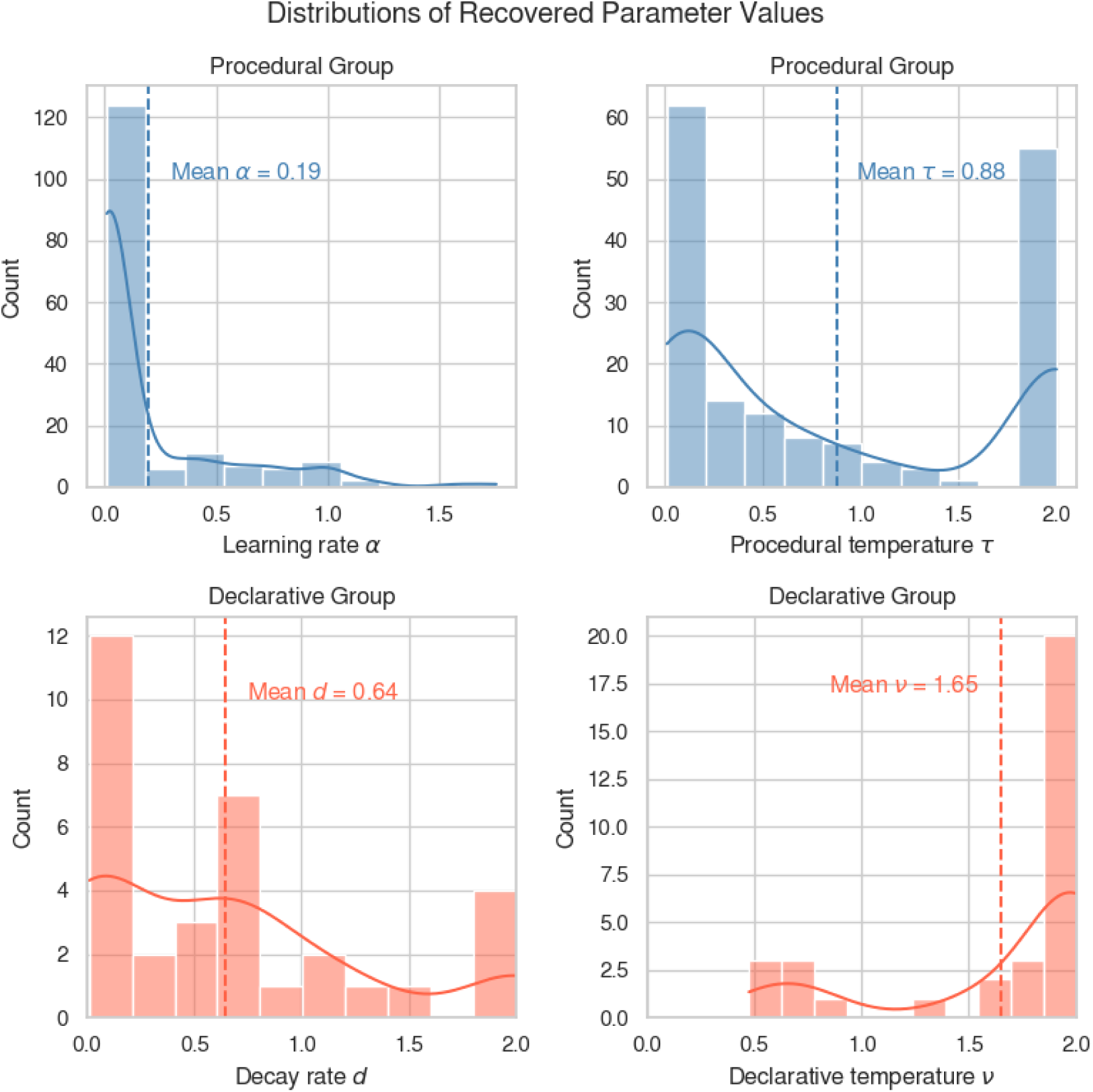
Histograms of best-fitting parameter values for individuals in the procedural (blue) and declarative (red) groups. Dashed vertical lines represent distribution means and solid lines represent kernel density estimates (kernel bandwidth = 0.75).

### Stability of Model Identification

Finally, we conducted a follow-up analysis to examine whether the process of model identification would be consistent over the course of the study. Of all 199 participants, 145 (72.9%; 135 in the procedural group and 10 in the declarative group) were best fit by the same model on both runs. Furthermore, the 54 individuals who were best fit by different models in the two runs exhibited very small differences between the log-likelihood of their best-fitting declarative and procedural models (Run 1: *M* = –2.41, *SD* = 13.84; Run 2: *M* = 1.71, SD = 3.71). This suggests that their behavior did not fall squarely within the predictions of a single model, and their different group assignments were more likely due to small behavioral changes rather than dramatic strategy switches. Altogether, these results suggest that the identification procedure is robust and that a participant’s tendency to make decisions in a procedural vs. declarative pattern is stable and does not fluctuate over the duration of an experiment.

### Group Differences In Behavioral Responses During The Task

Our prediction is that, because of the different dynamics in the two models, individuals best fitted by the declarative model would exhibit greater sensitivity to trial feedback and a greater probability of switching from their previous decision than individuals best fitted by the declarative model. To test this prediction, a dummy variable was created to code each trial after the very first as either a response switch (1) or not (0). The effect of the previous trial feedback on the probability of a response switch was then analyzed using a logistic mixed-effects model implemented in the lme4 package in R (Bates et al., 2015). The model included fixed effects of Group (declarative vs. procedural) and trial Feedback (Win vs. Loss vs. Neutral) as well as a random intercept and slope for each participant, having the form:

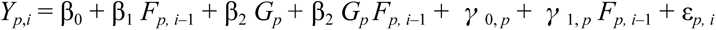

where *Y_p_*_,*i*_ is the *i*-th response (switch or not) of participant *p*, *G_p_*is the group assignment of participant *p*, *F_p,i_*_–1_ is the feedback received by participant *p* on trial *i*–1, and *γ*_0,*p*_ and *γ*_1,*p*_ are the individual intercepts and slopes of participant *p*.

The resulting linear model provided an acceptable fit to the data (Conditional *R*^2^ = 0.22). As predicted, we found a significant main effect of group (*β* = 1.10, *t* = 2.58, *p* < 0.001), a significant main effect of negative feedback (*β* = 1.45, *t* = 3.67., *p* < 0.001), and a significant interaction between the two factors (*β* = –1.12, *t* = –2.90, *p* = 0.003). As expected, although both groups were more likely to change their responses after a “loss” than a “win”, this difference was magnified in the declarative group (Figure 7, left). This finding was consistent with our understanding of the memory retrieval process, where more recent choices have a greater impact on the decision outcome..

**Figure 7.**
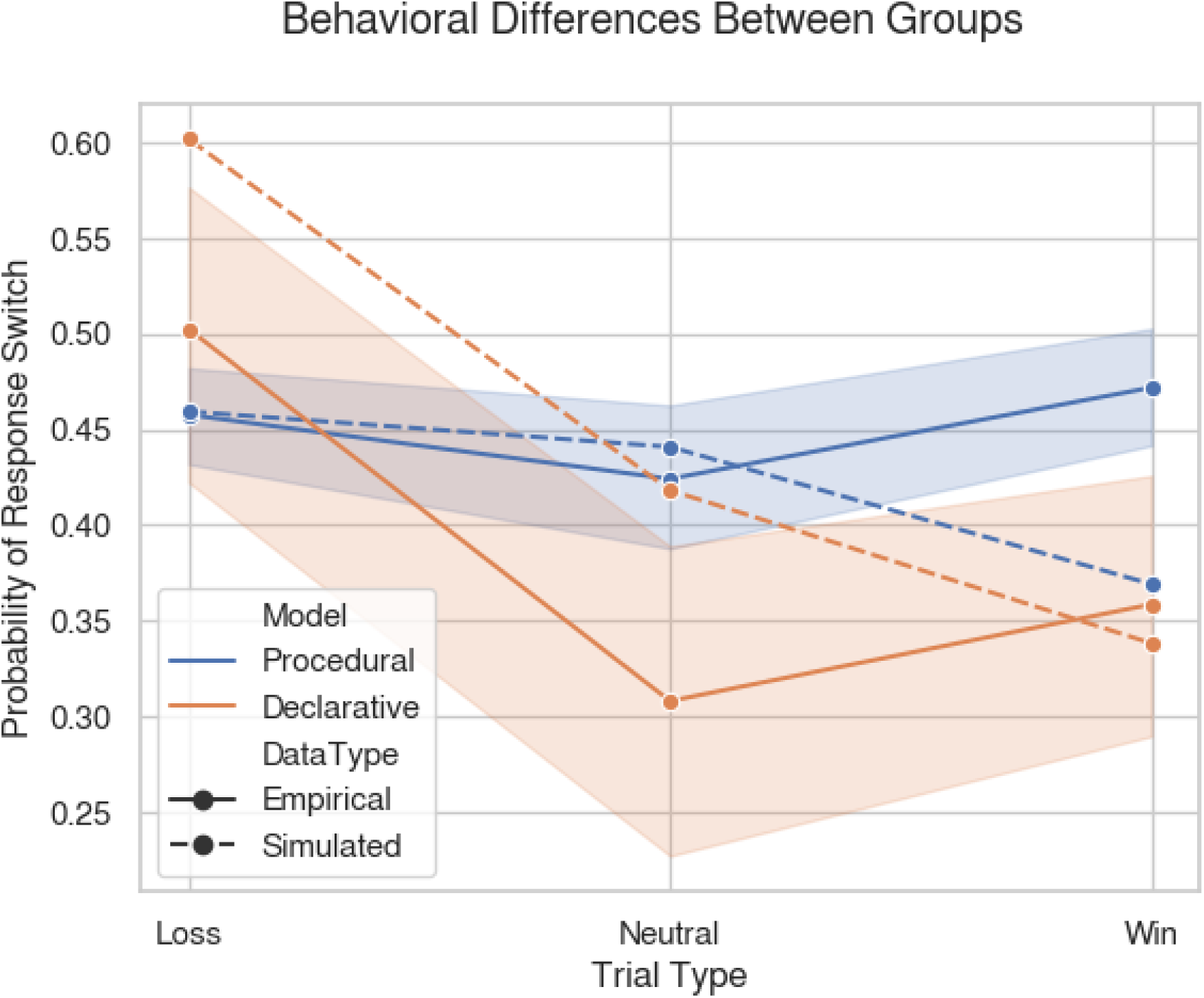
Main behavioral differences between participants best fit by procedural (blue) or declarative models (red). The data is consistent with the predicted pattern of behavior, with the effect of group and the effect of feedback interacting, and declarative participants exhibiting greater probabilities of response switches after a loss, but lower probabilities after a neutral or win feedback, than procedural ones. Continuous lines represent means from the empirical data; dashed lines represent means from forward simulations of the model; ribbons represent standard errors.

We also compared participants’ data against a new set of forward predictions. These predictions were generated by running the models with parameters sampled from the set of parameters values found in the participants best fit by each model. The results of this set of simulations are shown as the dashed lines of Figure 6. Note that these predictions were derived from models that were fit on the individual choices (“more” or “less”) made by participants, and not to the aggregated switching probabilities shown in Figure 6. Despite this, the models provided a remarkable fit to the data (mean squared error < 0.06). More importantly, the predictions of each model provide a better fit (a smaller mean squared error) to the corresponding group than to the other group.

### Group Differences in Brain Activity During the Task

In addition to finding significant behavioral differences between two groups of participants, we also found relevant and consistent differences in their task-based fMRI data. In line with previous studies using the same dataset (e.g., Delgado et al., 2000; Barch et al., 2013), our investigation focused on the difference in brain activity between “Mostly Win” and “Mostly Lose” blocks. This contrast is particularly important in this analysis, since participants are more likely to keep engaging their preferred decision-making system during “Mostly Win” blocks (as implied by Figure 5, right).

A *t*-test was performed to identify brain regions more active in one group over the other. The test was weighted using the relative log-likelihood of each participant’s being fitted by either of the two models, so that participants who had a stronger preference for one process were weighted more heavily (See Materials and Methods). The False Discovery Rate (Benjamini & Hochberg, 1995) procedure was used to correct for multiple comparisons at a level of *q* < 0.01. The analysis identified several brain regions that show significant BOLD signal differences between the procedural and declarative groups (Figure 6; Table 1). Specifically, participants in the procedural group (Figure 8, blue) showed greater task activation in the primary (inferior and superior occipital cortex and lingual gyrus) and secondary (left and right inferior temporal cortex) visual regions, and in the primary (M1) and secondary (supplementary motor area) motor regions, together with greater task activation in the bilateral insula. This pattern of activation is consistent with the acquisition of habitual stimulus-response associations.

**Figure 8.**
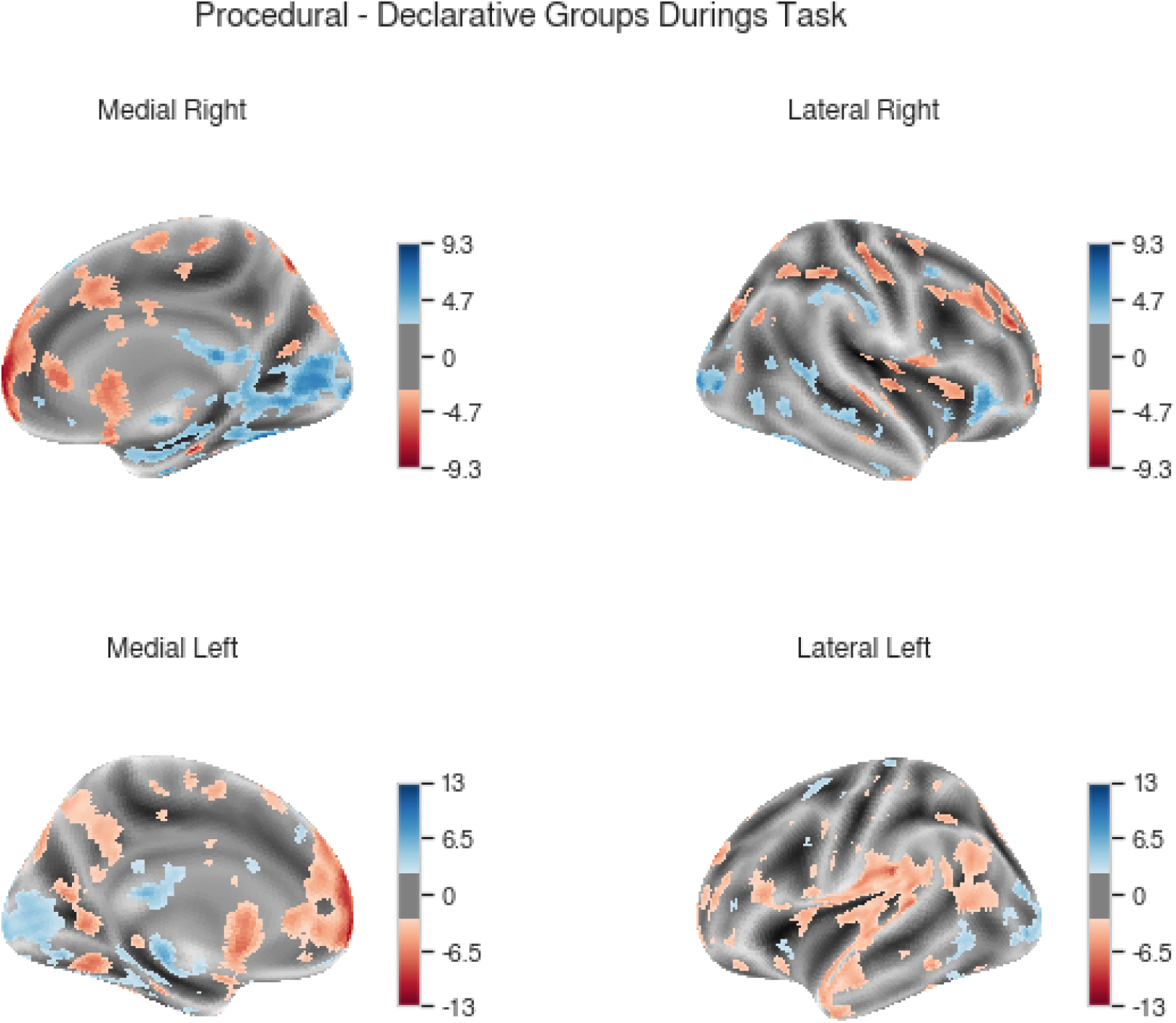
Regional brain activity differences between procedural and declarative participants during the incentive processing task. In the overlays, the color indicates the direction of the difference (blue: Procedural > Declarative; Red: Declarative > Procedural) and color intensity indicates the magnitude of the associated *T* statistics. Regions are thresholded at a significance level of at *q* < 0.01, corresponding to a voxel-wise threshold of *T* >= 2.70.

**Table 1:** Results from the task fMRI comparison. For clarity, only clusters spanning > 50 voxels and locations spanning > 1% of the cluster and > 10 voxels within the cluster are shown. L = Left, R = Right; the word “gyrus” is omitted from the location names.

| Contrast | MNI (Peak) | Automated Anatomical Labeling (AAL) Locations | Size (voxels) | Peak *T* value |
| --- | --- | --- | --- | --- |
| Procedural<br>><br>Declarative | 22, -82, -48 | R Cerebellum (4.44%); L Cerebellum (5.22%); R Fusiform (3.15%); L Calcarine (2.99%); R Calcarine (2.94%); R Lingual (2.76%); L Middle Occipital (2.64%); R Middle Temporal (2.14%); Left Fusiform (2.12%); L Lingual (1.62%); R Inferior Frontal (1.35%); R Inferior Temporal (1.27%); R Middle Occipital (1.21%); R Hippocampus (1.13%); L Inferior Occipital (1.12%); L Inferior Temporal (1.01%) | 30,662 | 14.09 |
|  | 62, -26, 46 | R Postcentral (34.73%); R Supramarginal (34.27%); R Inferior Parietal(17.60%); R Rolandic (11.89%) | 858 | 7.48 |
|  | -14, -34, 58 | L Paracentral (43.00%); L Postcentral (6.11%); L Precuneus (4.07%); L Precentral (1.53%) | 393 | 5.55 |
|  | -22, 4, 48 | L Superior Frontal (36.72%); L Middle Frontal (35.94%); L Supplementary Motor Area (3.52%) | 256 | 4.41 |
|  | -10, 36, 42 | L Superior Frontal (90.04%); ; L Middle Frontal (2.90%) | 241 | 5.23 |
|  | 36, 14, 60 | R Middle Frontal (96.94%); | 196 | 6.75 |
|  | 12, 24, 20 | R Anterior Cingulate (57.05%); | 149 | 5.49 |
|  | -6, 18, 26 | L Anterior Cingulate (40.74%); L Median Cingulate (10.37%) | 135 | 4.9 |
|  | -40, 16, -4 | L Insula (92.06%) | 126 | 5.1 |
|  | 10, -22, 66 | R Paracentral (43.93%); R Supplementary Motor Area (29.91%); R Precentral (26.17%) | 107 | 4.73 |
|  | -24, 68, 16 | L Middle Frontal (24.14%); L Superior Frontal (18.39%) | 87 | 4.61 |
|  | -40, -8, 52 | L Precentral (91.89%) | 74 | 4.15 |
|  | 28, 62, 24 | R Middle Frontal (56.16%); R Superior Frontal (39.73%) | 73 | 5.47 |
|  | -32, 0, 2 | L Putamen (56.72%) | 67 | 4.33 |
| Declarative<br>><br>Procedural | -4, 66, -2 | L Superior Frontal (7.26%); L Superior Temporal (3.29%); L Middle Temporal (3.25%); R Superior Frontal (5.56%); L Precuneus (2.12%); L Middle Frontal (2.12%); L Postcentral (2.00%); L Inferior Temporal gyrus (2.74%); L Inferior Frontal part (1.58%); L Angular (1.54%); L Rolandic (1.47%); L Anterior Cingulate (1.43%); L Supramarginal (1.35%); L Caudate (1.35%); R Median Cingulate (1.27%); R Superior Temporal (1.24%); L Precentral (1.17%); L Medial Frontal Cortex (1.13%); R Insula (1.10%); L Insula (1.06%); R Anterior Cingulate (1.05%); R Supplementary Motor Area (1.04%); L Temporal Pole (1.03%); L Supplementary Motor Area (1.01%); R Precuneus (0.99%) | 38,651 | -15.92 |
|  | 22, -44, -54 | R Cerebellum (86.87%) | 545 | -15.02 |
|  | 22, -24, -26 | R Parahippocampal (26.42%); R Cerebellum (33.97%); R Fusiform (18.24%) | 159 | -12.17 |
|  | 64, -4, 32 | R Inferior Frontal (38.31%); R Postcentral (31.17%); R Precentral (30.52%); | 154 | -9.32 |
|  | 40, 0, -44 | R Inferior Temporal (89.47%); R Temporal Pole (6.58%) | 152 | -7.13 |
|  | 2, -88, -16 | L Calcarine (25.49%); L Lingual (17.65%); L Crus (7.84%); | 102 | -8.99 |
|  | 6, -20, 76 | R Paracentral Lobule (59.60%); R Supplementary Motor Area (12.12%); | 99 | -7.52 |
|  | 52, -38, 56 | R Postcentral (89.53%) | 86 | -8.25 |
|  | 0, -4, 32 | R Median Cingulate (52.56%); L Median Cingulate (39.74%) | 78 | -4.21 |
|  | -42, 20, 50 | L Middle Frontal (98.68%) | 76 | -6 |
|  | 14, -26, -8 | R Lingual (15.71%); R Hippocampus (8.57%); | 70 | -4.72 |
|  | 40, 24, -26 | R Temporal Pole (98.33%); | 60 | -5.2 |

On the other hand, participants in the declarative group (Figure 7, red) showed greater brain activity in the canonical nodes of the default mode network (the bilateral medial frontal cortex, the bilateral precuneus, and the bilateral inferior parietal lobules) as well as regions involved in the encoding (right hippocampus, left anterior temporal lobe), and the retrieval (left and right lateral ventral prefrontal cortex) of declarative memories.

### Predicting Individual Differences in Decision-Making Processes Through Functional Connectivity

Functional connectivity data consisted of pairwise partial correlation matrices between each pair of the 264 regions in the Power parcellation scheme (Power et al., 2011). To identify which functional connectivity features predict whether a participant would belong to the declarative or procedural group, we used a logistic Least Absolute Shrinkage and Selection Operator (LASSO: Tibshirani, 1996) regression model, which reduces the number of potential predictors while retaining the most predictive features. The LASSO model was implemented using the *glmnet* package in R (Friedman et al., 2010). As noted in the Methods section, we used nested cross validation to ensure robust generalization of our results. This procedure yields multiple values of the hyperparameter λ, each of them estimated from a subset of the original data. We considered two possible values of λ: The median value and the value that yielded the greatest classification accuracy of the unseen data. Both values were then used to estimate the regression parameters, and their performance was tested again using canonical Leave-One-Out (LOO) cross-validation.

Table 2 and Figure 9 report the results of this procedure. The classification accuracy of the model was estimated at 91.1% for the median hyperparameter value (λ = 0.0059), and 100% when regressor parameters were re-estimated using the best hyperparameter value (λ = 0.0027).

**Figure 9:**
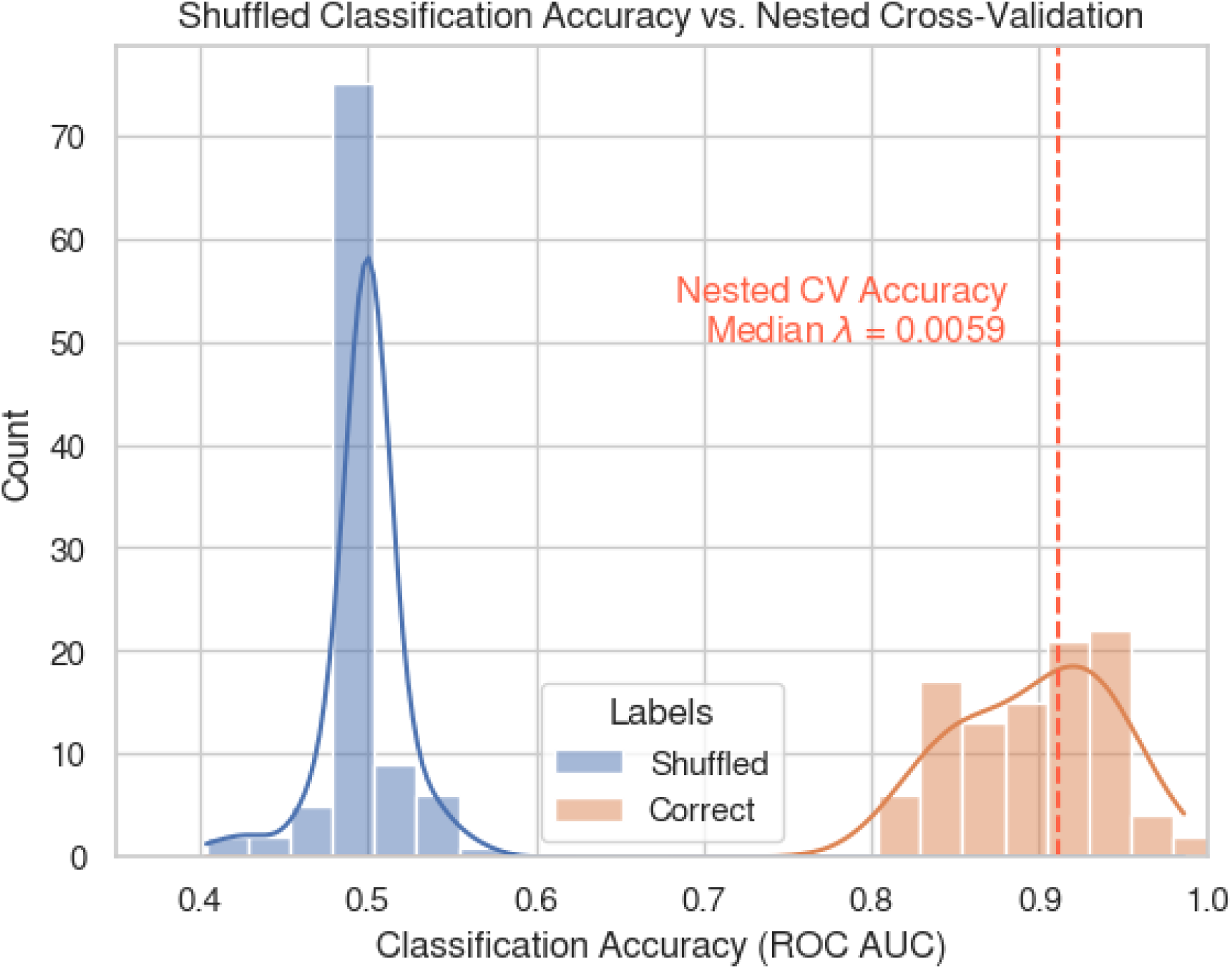
Results of the control classification analysis with correct (red) and randomly shuffled (blue) data. As expected, the classification accuracy of a LASSO model training on shuffled labels hovers at around 50%. Blue bars represent the number of simulations; solid lines represent the kernel density estimate (bandwidth = 1.5); the dashed line represents the accuracy of the model when the correct labels are applied and the median hyperparameter value is chosen.

**Table 2.**
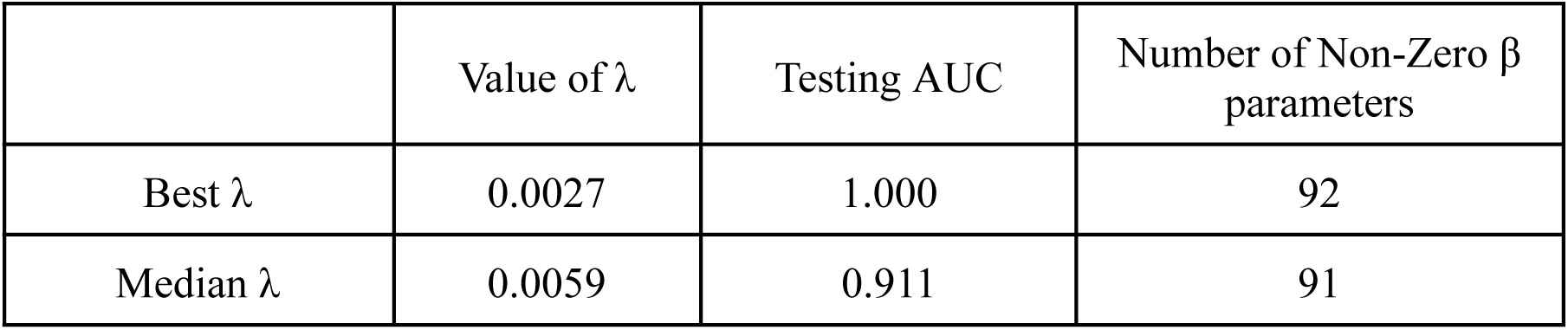
Summary of the LASSO Model Performance.

Although nested cross validation provides a robust safeguard against data leakage and other biases that could inflate accuracy rates, our results could still have been accidentally inflated by the upsampling of the declarative group and by the large number of features included in the regression. To ascertain that this was not the case, we conducted a bootstrap baseline comparison by re-running the analysis 100 times on the same data but with randomly shuffled labels. In each run, the LASSO model’s hyperparameter lambda was fitted using traditional LOO cross-validation (note that there is no reason to use nested cross-validation for randomly shuffled labels, as any bias and data leakage in traditional LOO would raise the baseline performance and thus provide a more stringent test of our hypothesis).

Figure 9 reports the distribution of ROC AUC scores obtained from each of these runs. As the figure shows, running the analysis with shuffled labels results in classification accuracies hovering around the chance level of 50%, and never exceeding 57%.

Finally, we examined which functional connectivity features were predictive of individual preferences in decision making. After LASSO regularization, 92 functional connections (approximately 0.11% of the total) had non-zero β parameters, suggesting a very sparse neurofunctional connectivity. Figure 10 shows the non-zero β weights of connections between networks and within networks as extracted from the fit LASSO model.

**Figure 10.**
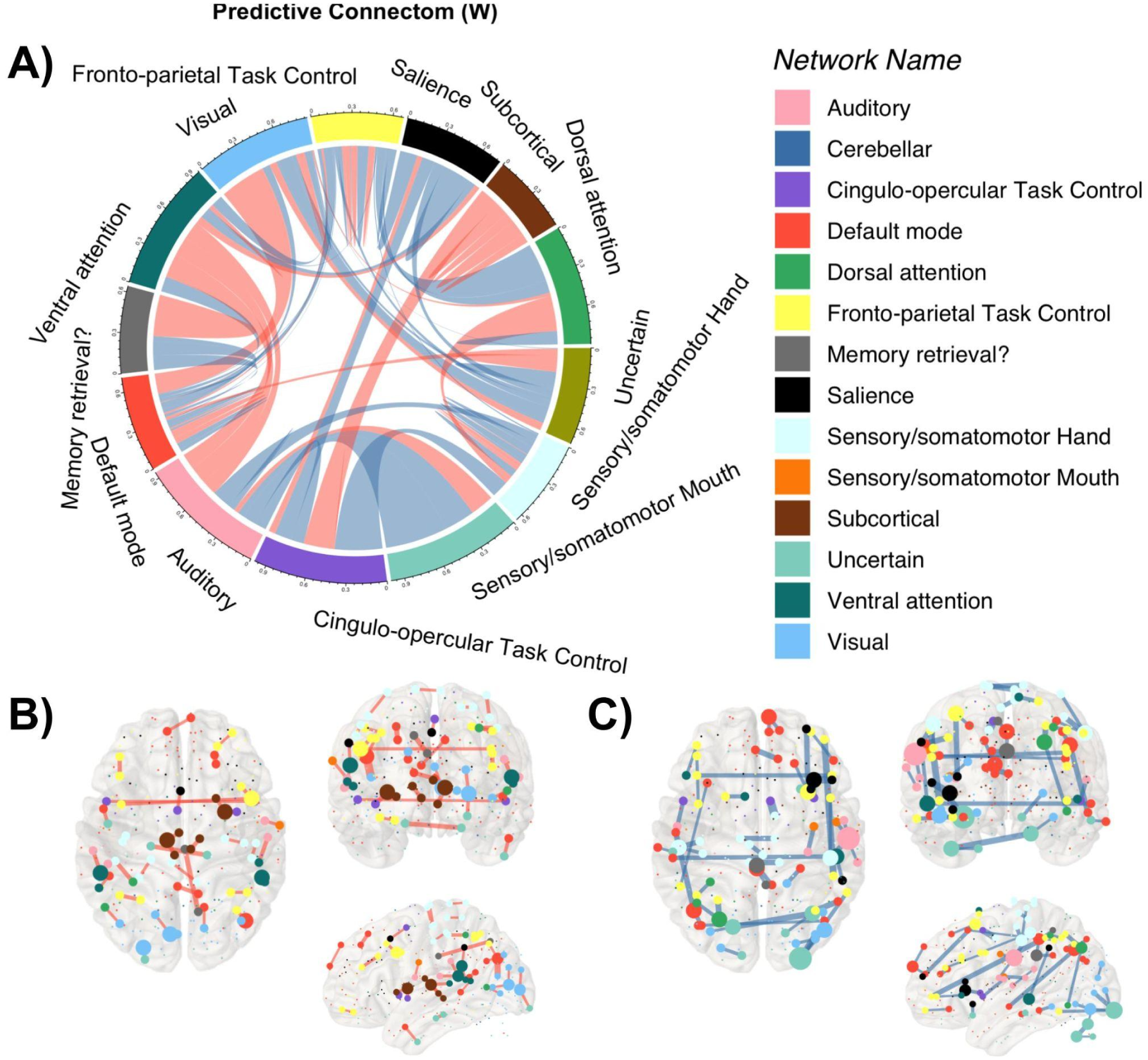
The distribution of predictive connections in declarative and procedural functional networks. (A) Predictive Connectom across the Power parcellation networks. Red connections denote positive weight values (*W* > 0), predicting the procedural Group, and Blue connections denote negative weights values (*W* < 0), predicting the procedural Group. (B) The anatomical location of the function connection is predictive of declarative group assignment; nodes represent regions, node colors indicate the network they belong to; node size indicates the degree of connectivity (the number of adjacent edges), and line width indicates the strength of connectivity between two nodes. (C) Anatomical location of the functional connections predictive of procedural group assignment.

Since the ultimate effect of a β parameter on the predicted group assignment depends on the polarity of the underlying functional connectivity, a positive β value has different implications if applied to a positive or negative partial correlation between two regions. Consider, for example, a connectivity value associated with a positive β value: if the underlying connectivity is positive (*r* > 0), then the degree of functional connectivity can be taken as a vote in favor of the declarative system. If, on the other hand, the underlying connectivity value is negative (*r* < 0), then it should be counted as a vote for the procedural system. To make the interpretation of the values unambiguous, we examined the corrected weight matrix by multiplying the β matrix with the sign of the functional connectivity matrix, obtaining a corrected weight correlation matrix *W*. The weight matrix *W* can now be interpreted unambiguously, since *W* > 0 predicts reliance on a declarative process and *W* < 0 predicts reliance on a procedural process. Fig 10A shows these different functional connections.

Connections predictive of declarative processes involve clearly different networks than connections predictive of procedural processes, as shown in Fig 10B and Fig 10C. As expected, reliance on declarative processes in decision-making was predicted by greater connectivity in the networks of regions associated with task control (task control networks and subcortical networks) as well as episodic memory (memory retrieval networks). Reliance on procedural learning processes, however, was predicted by greater connectivity in sensorimotor and salience regions. As Figures 10B-C show, the predictive functional connectivity also largely overlaps with the task-fMRI results, indicating that greater functional connectivity between two regions at rest predicts a greater likelihood that these two regions would be used to support the decision-making processes during the task.

## Discussion

This study shows that individuals rely on different mechanisms when deciding from experience, and that this preference is adaptive and reflects individual differences in the functional connectivity between each process’ corresponding circuitry. Specifically, individuals exhibiting stronger connectivity between and within frontoparietal and memory retrieval regions tend to use declarative strategies that are more reliant on episodic encoding and retrieval, while individuals with stronger connectivity in cingulate, sensory, and basal ganglia regions tend to rely on habitual actions and reinforcement learning. An individual’s preference can, in fact, successfully be predicted from their underlying functional connectivity.

Although our results shed new light on the neural basis of experiential decision-making, a number of limitations must be acknowledged. First, participants were assigned to the declarative or procedural group based on the log-likelihood of a corresponding cognitive model. Because no other ground-truth labels were available, these classifications should be interpreted with caution. Furthermore, the identification process resulted in two groups of different sizes, with most participants belonging to the Procedural group and only 17% to the Declarative group. This asymmetry is not uncommon in realistic datasets; there was no reason to expect an equal split of participants in the two groups and, in fact, previous studies of learning preferences found even more drastic differences in group sizes (Haile et al., 2021; 2023). Nonetheless, as acknowledged in the Results section, this asymmetry does create potential problems, and requires special care when applying machine learning methods.

Another limitation is that the incentive processing task differs from most decision-making paradigms, as there is no winning rule for participants to learn from the feedback. Thus, although this task was particularly well suited for the current study, more work is needed to determine whether these findings would translate to more realistic situations.

In addition, our study assumes that greater functional connectivity reflects greater efficiency. While much experimental evidence points to this, the mechanisms by which connectivity translates to computational efficiency are not clear. One proposed solution is that connectivity reflects better communication across regions. If so, connectivity should correspond to decision-making noise in our models. In fact, a follow-up analysis of individual model parameters shows that individual preferences for one process over the other largely follow differences in the estimated noise parameter. That is, individuals whose estimated procedural noise was larger than the estimated declarative noise were significantly more likely to be fit by the declarative model, and vice-versa (Welch Two Sample *T*-test: *t*(50.04) = 3.37, *p* = 0.001, see Figure 11).

**Figure 11.**
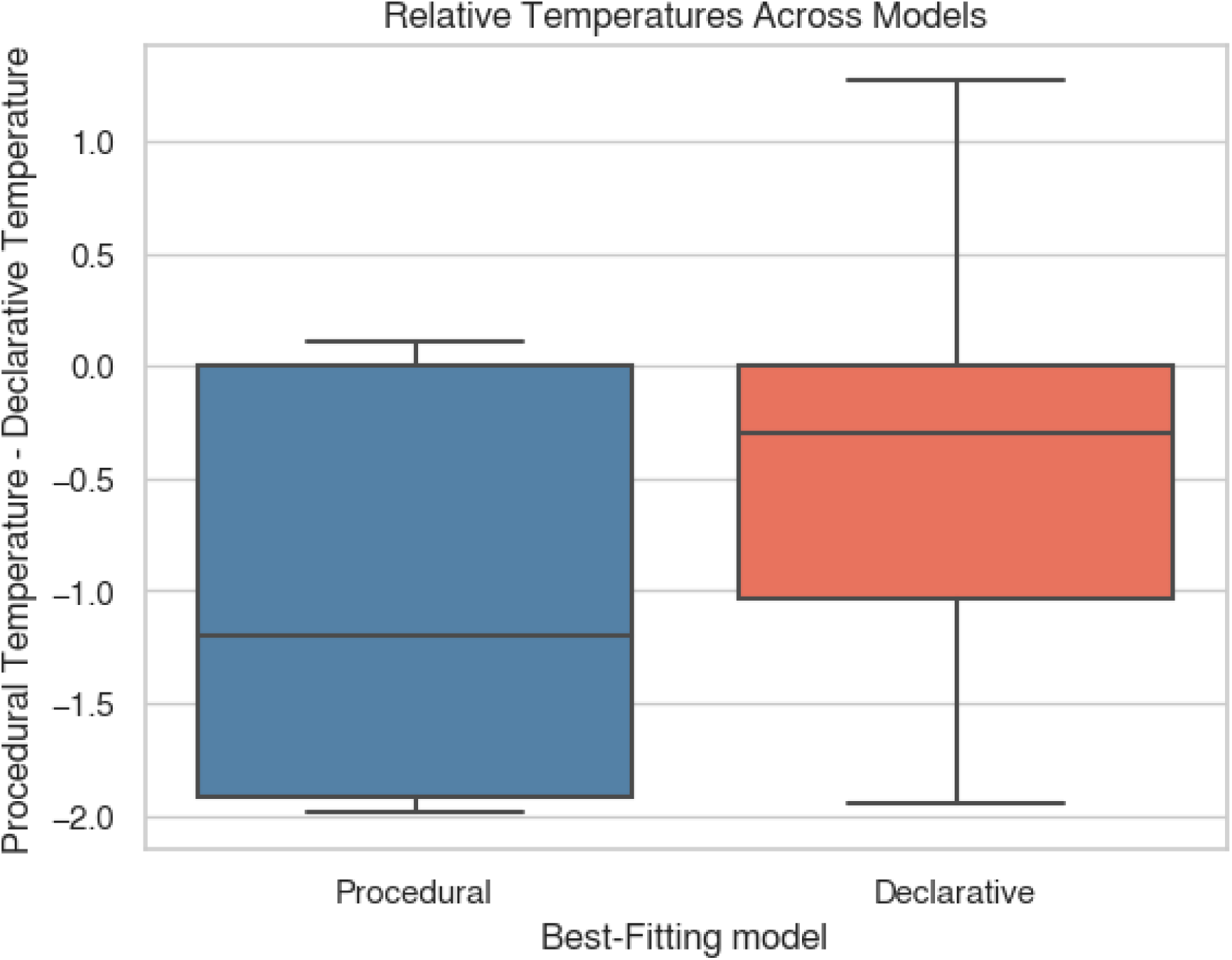
Differences in the estimated procedural and declarative temperature parameters τ and ν between the two groups. Individuals best fit by the procedural model had higher declarative temperature than individuals best fit by the declarative model, and vice-versa.

Although we have found evidence that individuals prefer to implement decision-making processes using the circuits that show the greatest connectivity, our analysis does not directly support a causal direction. To conclusively show a direct causal effect, it would be necessary to experimentally alter functional connectivity within the declarative and procedural circuits and measure subsequent changes in behavior in the same task. Such an experimental intervention is difficult to carry out in humans, although it is conceivable to use pharmacological interventions that directly target one circuit or intracranial direct stimulation of these circuits in patients with implanted electrodes or ECoG grids.

Finally, our study still does not directly address the source of differences in functional connectivity. Two hypotheses are possible. In principle, these differences could reflect underlying genetic or structural differences in brain function. Alternatively, it is possible that differences in functional connectivity simply reflect one’s past history of relying on one particular process (and its underlying circuit) over the other. In this case, individuals might have a preference (let’s say, for the declarative system), and repeated use leads to corresponding increases in functional connectivity (let’s say, in the memory retrieval network). The two explanations might be intertwined, i.e., small initial differences in the efficiency of these circuits might lead to greater use of one process over the other, and practice over time might result in additional gains in efficiency.

Further investigation is needed to fully untangle these questions, but these results nevertheless represent a promising step forward in directly connecting the mechanical underpinnings of brain circuitry with observed human behavior in decision-making.

## Data Availability Statement

All of the data and code are available in the OSF repository: https://osf.io/wf4my/

## Statements and Declarations

### Author Contributions

Conceptualization: AS, CS, YY. Methodology: AS,YY. Formal analysis: AS,YY. Visualization: AS,YY. Funding acquisition: AS. Project administration. AS; Supervision: AS. Writing – original draft: AS, YY, CS. Writing – review & editing: N/A

### Ethics approval

This study used data from the Human Connectome Project. The use of this data was approved by the University of Washington’s Institutional Review Board.

### Consent to participate

Consent to participate was acquired at the time the data was collected by the Human Connectome Project. The use of this data was approved by the University of Washington’s Institutional Review Board.

### Consent to publish

Participants consented to the publication of follow-up data analysis and results at the time their data was collected by the Human Connectome Project. The use of this data was approved by the University of Washington’s Institutional Review Board.

